# Subjective sleep onset latency is influenced by sleep structure and body heat loss in human subjects

**DOI:** 10.1101/2023.03.04.531123

**Authors:** Ryusei Iijima, Akari Kadooka, Kairi Sugawara, Momo Fushimi, Mizuki Hosoe, Sayaka Aritake-Okada

## Abstract

**Objectives:** The current study examined the relationship between subjective SOL, sleep structure, changes in skin and body temperature, and subjective evaluation in healthy young adults to elucidate the pathophysiological mechanisms of insomnia.

**Methods:** Twenty-eight participants (mean age: 21.5 ± 0.5 years) with no sleep problems participated in a 1-hour polysomnographic recording that obtained objective sleep parameters during the daytime while skin and body temperatures were recorded. The distal–proximal skin temperature gradient (DPG) was calculated. Subjective parameters, such as subjective SOL, sleep time, and restorative sleepiness, were evaluated before and after sleep.

**Results:** Most participants estimated their sleep latency as being longer than their actual SOL (13.7 min vs. 7.6 min). Objective SOL was significantly correlated with each sleep stage parameter whereas subjective SOL was negatively correlated with stage N2 sleep duration (Rho = −0.454, p = 0.020), slow-wave activity and delta power (Rho = −0.500, p = 0.011, Rho = −0.432, p = 0.031, respectively), and ΔDPG (the degree of reduction of heat loss before and after lights-off). Stepwise regression analysis showed that ΔDPG was the strongest predictive factor in explaining the length of subjective SOL.

**Conclusions:** The degree of heat dissipation before falling asleep contributed most to the sensation of falling asleep in healthy young adults. This finding may be helpful for elucidating the physiological mechanisms of insomnia and its treatment.

**STATEMENT OF SIGNIFICANCE:** The time estimate ability is also activated during sleep. However, some insomniacs have abnormalities in this function and suffer from sleep state misperception, which is a discrepancy between subjective and objective sleep time. They often overestimate the sleep onset latency. We investigated the relationship between subjective sleep onset latency, sleep structure, EEG frequency components, and body temperature during the process of falling asleep in healthy adults. It was suggested that the Stage N2 duration, the amounts of slow wave activity around after lights-out, and the degree of heat dissipation before falling asleep may be related to the perception of falling asleep. These results are expected to contribute to the understanding of the pathophysiological mechanisms and to the treatment of insomnia.

## INTRODUCTION

Several previous studies reported that humans possess time estimation ability (TEA) not only during wakefulness, but also during sleep [1,2]. Previous studies of TEA during sleep reported a relationship between subjective sleep duration and objective sleep duration, the endocrine system, the autonomic nervous system, the amount of daytime activity, and psychological factors [3–6]. Our previous study revealed that TER, as an indicator of a subjectively-estimated time interval, was higher at the beginning of the sleep period (i.e., sleep time was overestimated compared with the actual time elapsed), and that it successively decreased toward the end of sleep [1]. Additionally, the study reported that the greater the amount of SWS obtained by the study participants, the longer the sleep time they subjectively experienced, whereas that greater the amount of rapid eye movement (REM) sleep the study participants obtained, the shorter the sleep time they subjectively experienced.

In recent years, various findings in patients with insomnia have indicated abnormalities in TEA [7]. It has been shown that patients with insomnia, such as primary insomnia and psychophysiological insomnia, tend to rate their sleep duration as shorter compared with healthy participants [8]. Sleep state misperception (International Classification of Sleep Disorders: ICSD1997, 2nd ICSD, 3rd ICSD) has been reported to show a particularly clear discrepancy between subjective and objective sleep duration in insomnia [9]. It has been reported that patients with sleep state misperception not only significantly underestimate the duration of sleep they receive, but also overestimate sleep onset latency (SOL) [10–12]. To date, many studies examining the pathophysiology and background factors of insomnia and paradoxical insomnia have been conducted using electroencephalography (EEG) and brain imaging methods, as well as psychological studies such as sleep stages focused on sleep duration, brain imaging, quantitative EEG analysis, and cyclic alternative pattern analysis [13–17]. The results revealed that the amplitudes of alpha power, sigma power, and beta power in fast Fourier transform (FFT) analyses during non-REM sleep [18] as well as alpha power and sigma power during REM sleep [19] in patients with sleep state misperception were significantly higher than those in other insomnia patients.

However, most previous studies of insomnia have focused on “sleep duration” during the entire night or EEG components during sleep duration, whereas very few studies have focused on subjective sleep latency, which is one of the crucial characteristics of sleep state misperception. One study reported that patients who misidentified SOL exhibited more instances of waking after sleep onset and required longer periods of uninterrupted sleep to recognize that they had fallen asleep, compared with healthy participants [20].

The process of falling asleep is one of the most important aspects of timing for determining individuals’ subjective evaluation of sleep, as well as changes in physiological function, such as brain system and body temperature [21–23]. Heat dissipation can be measured indirectly by assessing the temperature gradient between the distal (dorsal hands and feet) and proximal (subclavian, forehead) skin temperatures, which is known as the distal-proximal skin temperature gradient (DPG). The DPG is reported to be a good predictor of sleepiness, and is correlated with SOL [23,24]. Nevertheless, few studies have focused on the relationship between the characteristics of subjective SOL and changes in temperature and sleep structure, which are closely related to sleep onset. Clarifying the relationship between TEA, body temperature, and sleep structure during the process of falling asleep may provide helpful clues for elucidating the pathophysiological mechanisms of insomnia patients.

The objective of the current study was to examine the relationship between subjective SOL, sleep structure, changes in body temperature (skin temperature and tympanic membrane temperature), and subjective evaluation in healthy young adults as a infrastructural study to help elucidate the pathophysiological mechanisms of insomnia.

## METHODS

### Participants

This study was approved by the Ethics Committee of Saitama Prefectural University (Approval No. 30055, 19024). Participants were 28 healthy adult volunteers (male: 7, female: 21, mean age: 21.5 ± 0.51 years). All participants provided written, informed consent after receiving an explanation of the possible risks and details of the study. The inclusion criteria were as follows: 20–40 years of age; regular sleep habits. Participants were instructed to maintain a regular sleep-wake schedule, record their sleep patterns in a sleep log, and wore a wrist-activity recorder (Active style Pro, OMRON co. Kyoto, Japan. or ActiGraphLink, Acti Japan co. Chiba, Japan) for 1 week before the experiment. Habitual sleep-onset and sleep-offset times were confirmed using these records. The exclusion criteria were as follows: history of sleep-related or psychiatric disorders; neurological, psychiatric, or sleep disorders; history of taking psychoactive drugs, antihistamines, medications for cardiovascular disease for 1 week before the experiment; other chemicals with anti-hypnotic effects (caffeine, nicotine-containing products, or alcohol) for 24 h before the experiment; shift-work; and travel to a different time zone within 1 month of the experiment. Female participants participated only in the follicular period because of the effects of the menstrual cycle on sleep and body temperature. In addition, the Pittsburgh Sleep Quality Index (PSQI) [25], Morningness - Eveningness Questionnaire (M ·E) [26], Epworth Sleepiness Scale (ESS) [27], Medical Outcome Study (MOS) 36-Item Short-Form Health Survey (SF-36) [28], The General Health Questionnaire (GHQ) [29], and the Center for Epidemiologic Studies Depression Scale (CES-D) [30] were conducted as pre-screening measures to confirm the absence of sleep disorders and psychiatric disorders prior to the experiment.

### Experimental procedure

The experimental schedule is shown in Figure 1. All experiments were performed in an isolated shield room at the laboratory of Saitama prefectural university in Japan. Participants arrived at the laboratory at 13:00 and were instructed to go to bed at 14:00 (lights-off). Polysomnography (PSG) during the daytime was carried out after electrodes and sensors were attached, and participants were woken up at 15:00 (lights-on).

**Fig 1.**
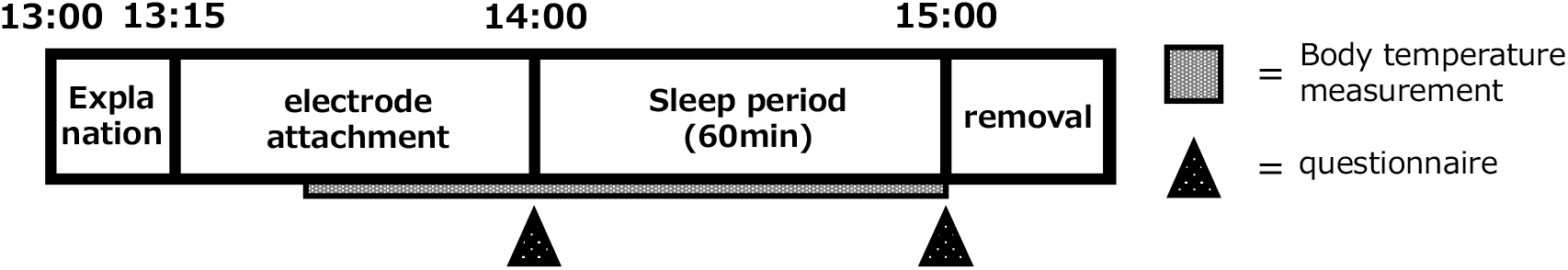
The experimental schedule

Tympanic membrane temperature as a measure of core body temperature (CBT) was recorded every 1 min throughout the experimental period using a portable body temperature logger (resolution 0.02 °C; LT-8; Gram Corporation, Saitama, Japan). An ear dedicated probe for tympanic membrane temperature was self-inserted into the ear. Additionally, distal (foot and hand) and proximal (forehead, subclavian) temperatures were recorded every 1 min throughout the experimental period using button-typed skin temperature sensors (resolution 0.1 °C; Thermocron SL type, KN Laboratories, Inc.). Skin temperature probes were covered by thermal-insulating material to minimize differences in skin temperature and kept in place with surgical tape.

PSG comprised EEG (Fp1-A2, Fp2-A1, F3-A2, F4-A2, C3-A2, C4-A1, O1-A2, and O2-A1) in accordance with the 10–20 electrode system, electro-oculography (left-A2 and right-A1), chin surface electromyography, and electrocardiography. PSG data were obtained continuously during each experiment, digitalized at 500 Hz, and stored in a digital EEG system (EEG1200, Nihon Kohden, Tokyo, Japan). High- and low-cut filters were set at 35 Hz (except chin surface electromyography at 100 Hz) and 0.3 Hz, respectively.

Subjective parameters, such as subjective SOL, and sleep duration were evaluated after the sleep period. Other subjective parameters such as sleepiness, mood, physical and mental fatigue, and restorative sleepiness (only after sleep period) were evaluated using a visual analog scale (VAS) and Stanford Sleepiness Scale scores [31] at bedtime and after waking, respectively. Throughout the experiment, room temperature (23°C), and illuminance (150 lx during wakefulness; 0 lx during sleep) were regulated. All participants used the same bedding (mattress, pillow, and comforter). Meals were controlled to ensure a standard daily calorie intake for all participants. Participants were allowed to drink only water, and the quantity was not limited. Participants were prohibited from sleeping during the experiment until the scheduled bedtime and were monitored by researchers. Participants were also prohibited from the consumption of caffeine or alcohol on the day before the experiment.

## DATA ANALYSES

### Sleep EEG parameters

PSG data obtained 1 h were scored for 30-s epoch periods, according to the standard criteria of the American Academy of Sleep Medicine, version 2.6 [32], by two scorers (a certified scorer (registered polysomnographic technologist) rescored and double-checked after one scorer scored) who were blinded to each participant’s condition (baseline or exercise) using the G3 (Philips Japan co, Tokyo, Japan). The lengths of stage W, stage N1, stage N2, and stage N3 (SWS) and REM sleep, sleep latency, wake after sleep onset, total sleep time, sleep period time, sleep efficiency, and arousal index for the total sleep time were calculated for all PSG recordings. The frequency of body movement during sleep was determined by monitor viewing and EEG waveforms in real time. For a more detailed analysis of sleep structures, power spectral analysis using the FFT algorithm, a parameter of brain activity at the cortical level, was also conducted with CSA Play Night Owl Professional (NoruPro Light Systems, Tokyo). The EEG power values during non-REM sleep (stages N2 and N3) were obtained for C3-A2 and C4-A1 using FFT analysis with 2,048 data points (eight, 4.096-s data sequences), tapered by the Hanning window: slow-wave activity (SWA): 0.5–2 Hz, delta (δ): 0.5–4.0 Hz, theta (θ): 4–8 Hz, alpha (α): 8–13 Hz, slow sigma (σ): 10–13 Hz, fast sigma (σ): 13–16 Hz, and beta (β): 13 Hz. Because the eight 4.096-s sequences were longer than the visual epoch definitions for sleep staging, both sides of each sequence overlapped by 0.39 s. The eight FFT values for each of the eight sequences were summed to obtain the value of EEG power per epoch. Epochs containing artifacts were visually identified and excluded from the analysis.

### Body temperature parameters

During the experiment, tympanic membrane temperature and distal and proximal skin temperature for the calculation of DPG were recorded every 1 min, and telemetric signals were stored in a computerized monitoring system, then transferred to a personal computer. Mean CBT and distal and proximal skin temperature data were averaged for every 5-min period during daytime sleep for 1 h after lights-off, and every 5 min for the 15 min before lights-off. Distal-proximal skin-temperature gradient (DPG), which is one of the parameter for evaluating sleep onset, was calculated from the difference between the distal skin temperature and the proximal skin temperature. To compare the changes in DPG before and after bedtime, ΔDPG (the degree of reduction of heat loss) was calculated by subtracting the value of DPG during the 15 min after lights-off from the value of DPG during the 15 min before lights-off [33].

### Statistical analysis

Two of the 28 participants were excluded from the analysis because their subjective SOL or objective SOL values significantly deviated from the mean values (> ± 2 standard deviations).

The Shapiro-Wilk test was used to determine whether the data sets were normally distributed. Spearman’s correlation coefficient values of the subjective sleep parameters and objective sleep parameters, and of the body temperature before and after lights-off, were also evaluated. SPSS version 26.0 (IBM SPSS, Armonk, NY) was used for all statistical analyses. Results are presented as means and standard error values, except for the parameters shown in the tables. The level of significance was set at P < 0.05. Stepwise regression analysis was conducted for the prediction of the subjective SOL.

## RESULTS

### Demographic data and sleep parameters of the participants

Table 1 shows the demographic data, subjective SOL, and objective sleep parameters obtained from the sleep EEG analysis of 26 participants. The total sleep time was 43.29 ± 16.66 min, and sleep efficiency was 72.13 ± 27.80%. The subjective SOL was 13.70 ± 8.10 min, and the objective SOL was 7.57 ± 8.80 min. Although the SOL and PSQI values were low, it is unlikely that morbid sleepiness was the cause.

**Table 1.** General characteristics and sleep parameters

|  | Mean±SD, N(%) |
| --- | --- |
| N | 26 |
| Age(year) | 21.54±0.50 |
| Male | 7(26.92) |
| BMI | 20.64±2.44 |
| PSQI | 5.96±2.10 |
| Subjective SOL(min) | 13.70±8.10 |
| Objective SOL(min) | 7.57±8.80 |
| Total sleep time(min) | 43.29±16.66 |
| Wake time after sleep onset(min) | 8.51±11.14 |
| Sleep efficiency(%) | 72.13±27.80 |
| Stage N1(min) | 10.74±6.97 |
| Stage N2(min) | 21.30±10.43 |
| Stage N3(min) | 11.28±11.85 |
| Stage N1 latency(min) | 6.17±7.89 |
| Stage N2 latency(min) | 9.85±8.69 |
| Stage N3 latency(min) | 25.48±11.04 |

### Relationship between subjective SOL and objective SOL

The values for the difference between subjective and objective SOL for all participants are shown in Figure 2a. Most participants overestimated subjective SOL, which was longer than objective SOL. Moreover, no significant correlation was found between subjective SOL and objective SOL (Rho = 0.345, p = 0.085) (Figure 2b).

**Fig 2.**
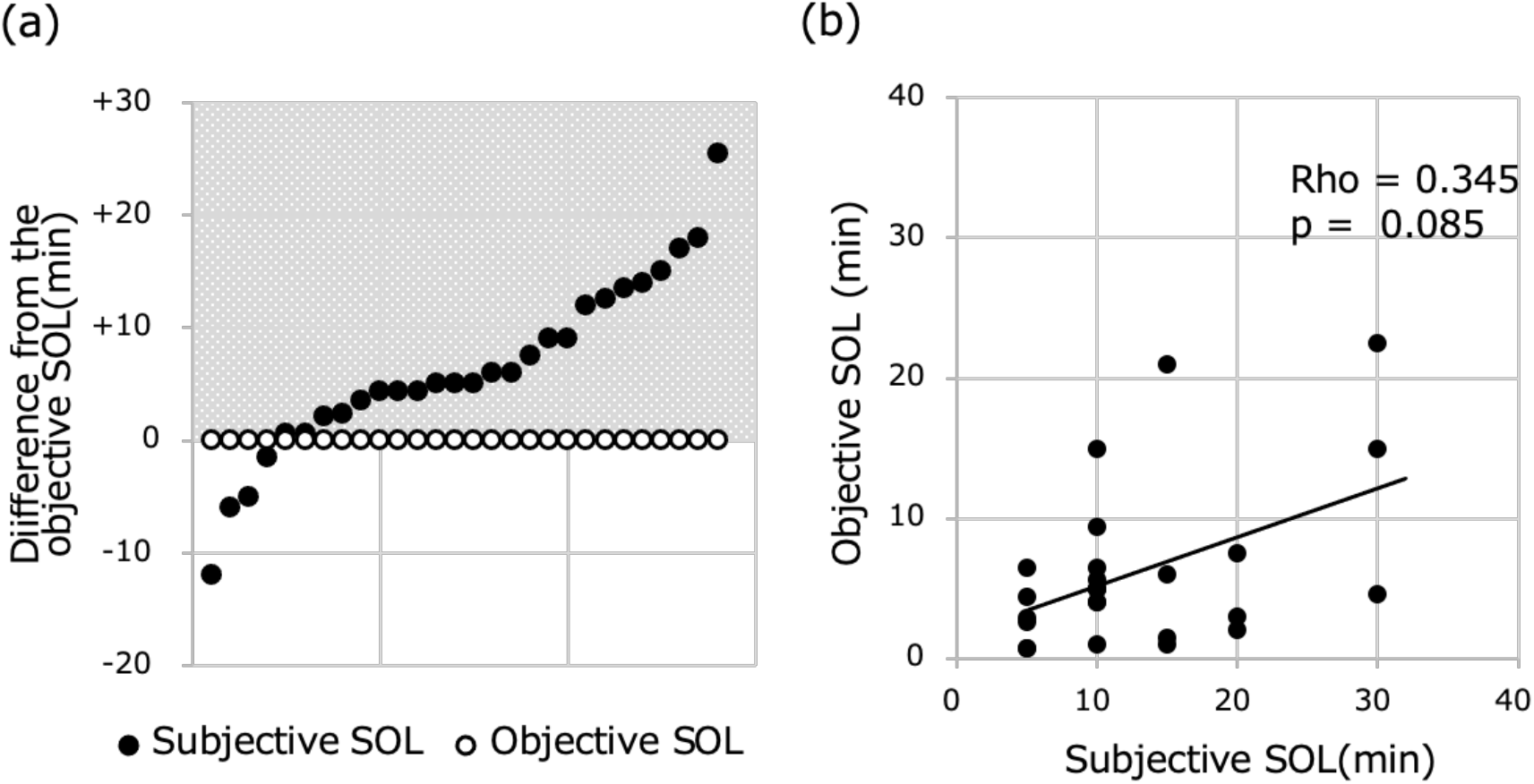
The vertical axis indicates the difference from the objective SOL, and the black markers indicate the subjective SOL (a). The shadow background area indicates that the SOL was estimated to be long. Relationship between subjective SOL and objective SOL (b).

### Relationship between each SOL and objective sleep parameters

The correlations between objective SOL, subjective SOL and the latency of each sleep stage are shown in Figure 3. Objective SOL showed a significant positive correlation with stage N1 latency (Rho = 0.963, p < 0.001), stage N2 latency (Rho = 0.922, p < 0.001), and stage N3 latency (Rho = 0.626, p = 0.004), respectively (Figure 3a, 3b, 3c). On the other hand, subjective SOL was not significantly correlated with stage N1 latency (Rho = 0.347, p = 0.082), stage N2 latency (Rho = 0.402, p = 0.051), and stage N3 latency (Rho = 0.389, p = 0.099), respectively (Figure 3d, 3e, 3f).

**Fig 3.**
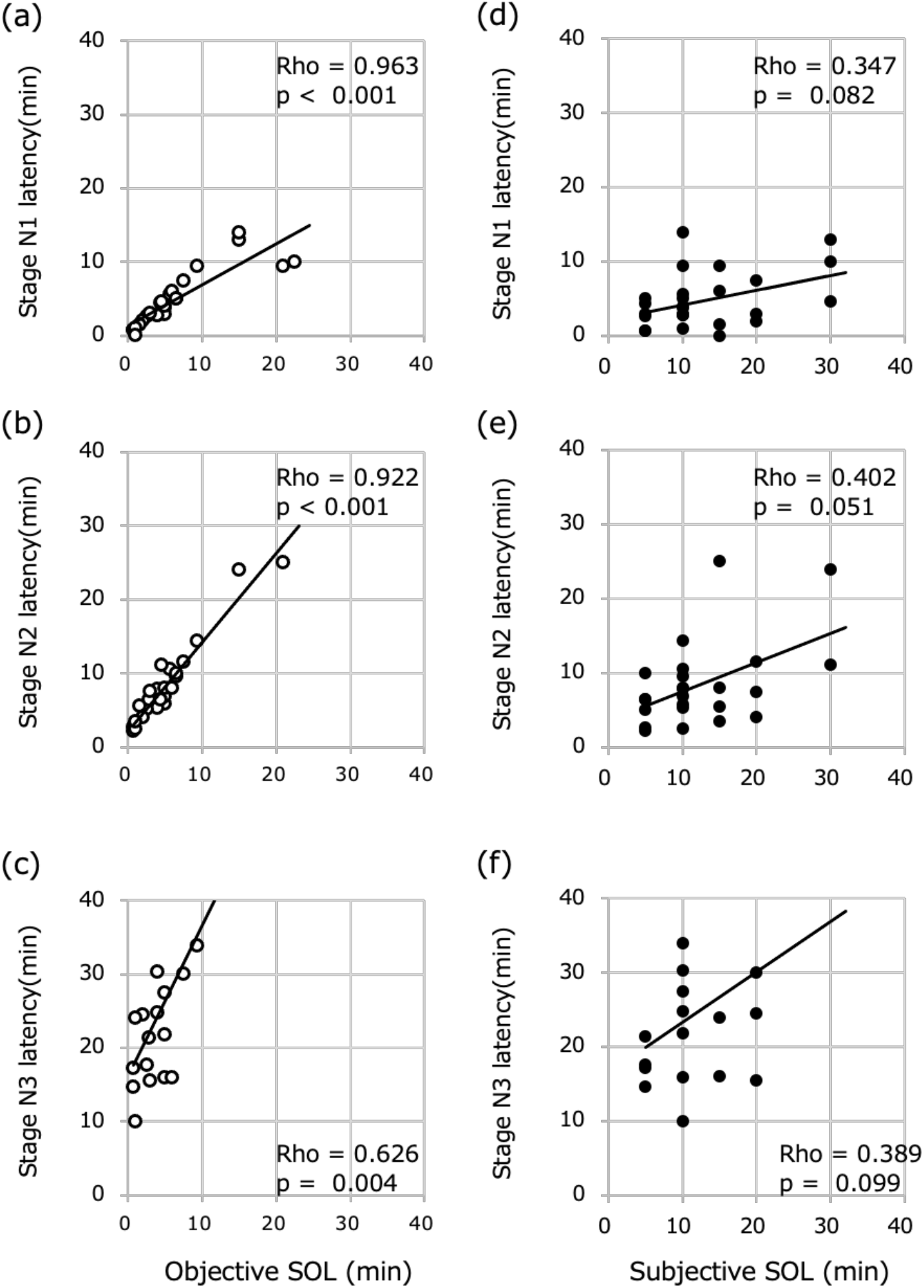
Relationships between objective SOL and latency of sleep stages: (a) stage N1 latency, (b) stage N2 latency and (c) stage N3 latency. Relationships between subjective SOL and latency of sleep stages: (d) stage N1 latency, (e) stage N2 latency and (f) stage N3 latency.

The correlations between objective SOL, subjective SOL and the duration of each sleep stage are shown in Figure 4. Objective SOL showed a significant positive correlation with stage N1 duration (Rho = 0.514, p = 0.009) and a significant negative correlation with stage N3 duration (Rho = −0.633, p < 0.001). However, objective SOL did not show a significant correlation with stage N2 duration (Rho = −0.368, p = 0.065). Subjective SOL showed a significant negative correlation only with stage N2 duration (Rho = −0.454, p = 0.020), and no significant correlation with stage N1 (Rho = 0.172, p = 0.412) or stage N3 duration (Rho = −0.023, p = 0.911).

**Fig 4.**
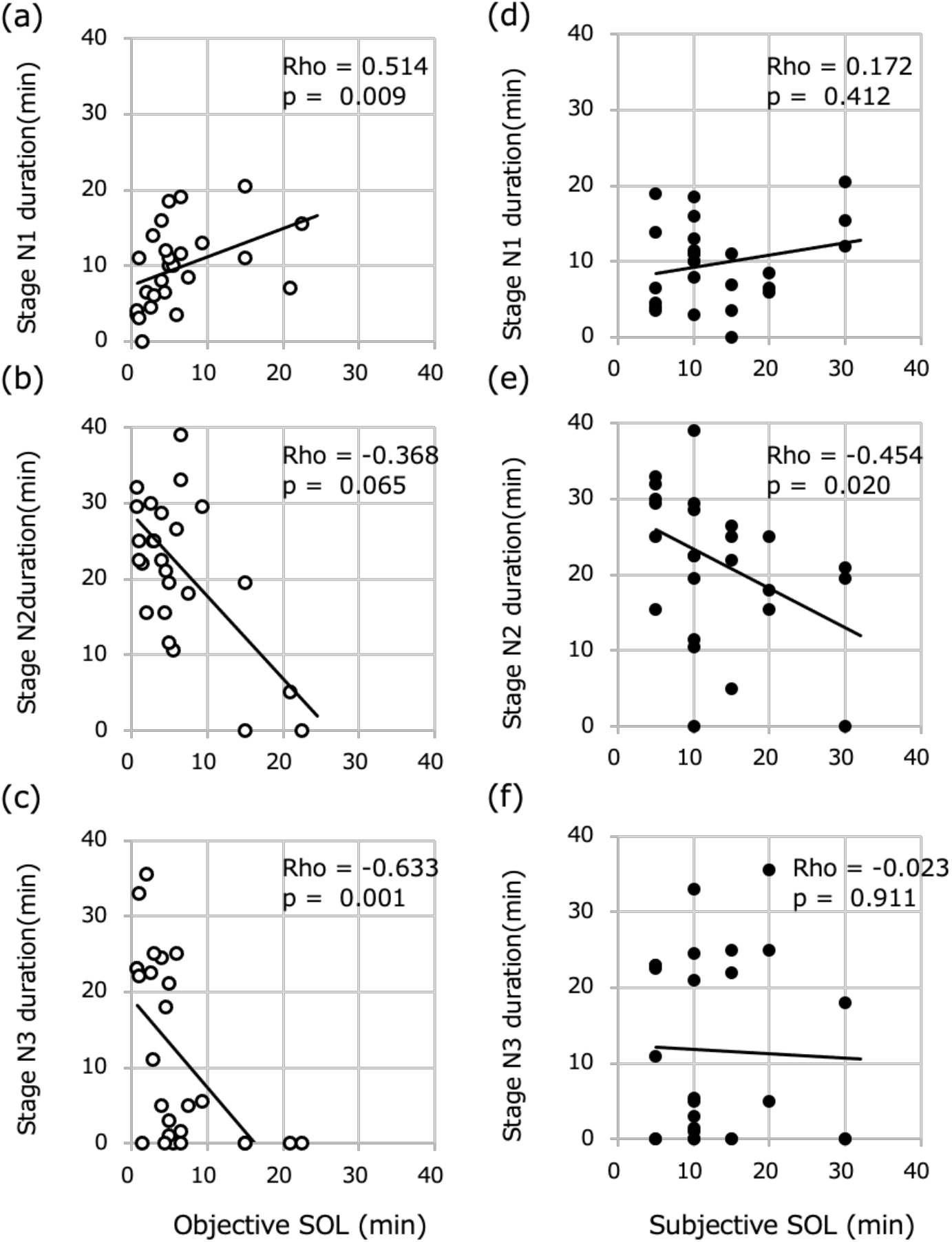
Relationships between objective SOL and duration of sleep stages: (a) stage N1 duration, (b) stage N2 duration and (c) stage N3 duration. Relationships between subjective SOL and duration of sleep stages: (d) stage N1 duration, (e) stage N2 duration and (f) stage N3 duration.

### Relationship between each SOL and the power of each frequency

Table 2 shows the average power of each frequency during sleep in the 60 min after lights-off and during sleep in the 15 min after the lights-off, and their correlation with SOL. Of the 26 participants, data from one participant were excluded from the analysis for the 15 min after lights-off because they did not exhibit a sleep epoch in the period. Objective SOL showed a significant negative correlation with SWA, δ, θ, and σ (p < 0.05 for each; p-values shown in the table) in the 60 min period after lights-off, while subjective SOL showed no significant correlation with the power values of each frequency. Regarding the relationship between the power of each frequency during sleep in the 15 min after lights-off and SOL, objective SOL showed a significant negative correlation with the power of all frequencies in the 15 min period after lights-off (p < 0.05 for each, p-values are listed in the table). Subjective SOL showed significant negative correlations with SWA (Rho = −0.500, p = 0.011) and δ power (Rho = −0.432, p = 0.031) (Table 2). There was no significant correlation between the power of each frequency and SOL in the second, third, and fourth 15-min sleep periods.

**Table 2.** The average power of each frequency during sleep in the 60 minutes after the lights-off and during sleep in the 15 minutes after the lights-off, and their correlation with sleep onset latency

| Power of 60 minutes after<br>lights-off (n=26) |  | Objective SOL |  | Subjective SOL |  |
| --- | --- | --- | --- | --- | --- |
|  | mean±SD | rho | p | rho | p |
| SWA | 16371.0±8290.0 | −0.663 | 0.000* | −0.200 | 0.327 |
| δ | 28345.2±12874.8 | −0.629 | 0.001* | −0.232 | 0.253 |
| θ | 15766.4±4809.3 | −0.440 | 0.025* | −0.109 | 0.595 |
| α | 12141.5±4014.4 | −0.388 | 0.050 | −0.023 | 0.911 |
| β | 12104.9±3107.3 | −0.313 | 0.119 | −0.071 | 0.730 |
| σ | 7350.4±2280.1 | −0.400 | 0.043* | −0.032 | 0.875 |
| slow σ | 7138.6±2545.3 | −0.389 | 0.050 | −0.020 | 0.924 |
| fast σ | 5377.3±1582.3 | −0.381 | 0.055 | −0.055 | 0.788 |
| Power of 15 minutes after<br>lights-off (n=25) |  |  |  |  |  |
| SWA | 9059.3±3311.3 | −0.699 | 0.000* | −0.500 | 0.011* |
| δ | 18213.9±6414.7 | −0.661 | 0.000* | −0.432 | 0.031* |
| θ | 13446.0±3805.7 | −0.515 | 0.008* | −0.284 | 0.169 |
| α | 10140.9±2972.6 | −0.465 | 0.019* | −0.255 | 0.219 |
| β | 12125.2±3031.5 | −0.500 | 0.011* | −0.200 | 0.337 |
| σ | 6573.8±1999.4 | −0.616 | 0.001* | −0.235 | 0.257 |
| slow σ | 5908.9±1814.0 | −0.505 | 0.010* | −0.255 | 0.219 |
| fast σ | 4983.6±1511.0 | −0.628 | 0.001* | −0.233 | 0.262 |

### Relationship between SOL and DPG

Objective SOL was not significantly correlated with mean DPG during 15 min before and after lights-off (before lights-off: Rho = −0.193, p = 0.344; after lights-off: Rho = −0.179, p = 0.405). Additionally, subjective SOL was not significantly correlated with mean DPG during 15 min before and after lights-off (before lights-off: Rho = −0.279, p = 0.168; after lights-off: Rho = −0.037, p = 0.859). In contrast, subjective SOL showed a significant negative correlation with ΔDPG (Rho = −0.639, p < 0.001) (Fig. 5). No significant correlation was found between objective SOL and ΔDPG (Rho = −0.097, p = 0.637).

**Fig 5.**
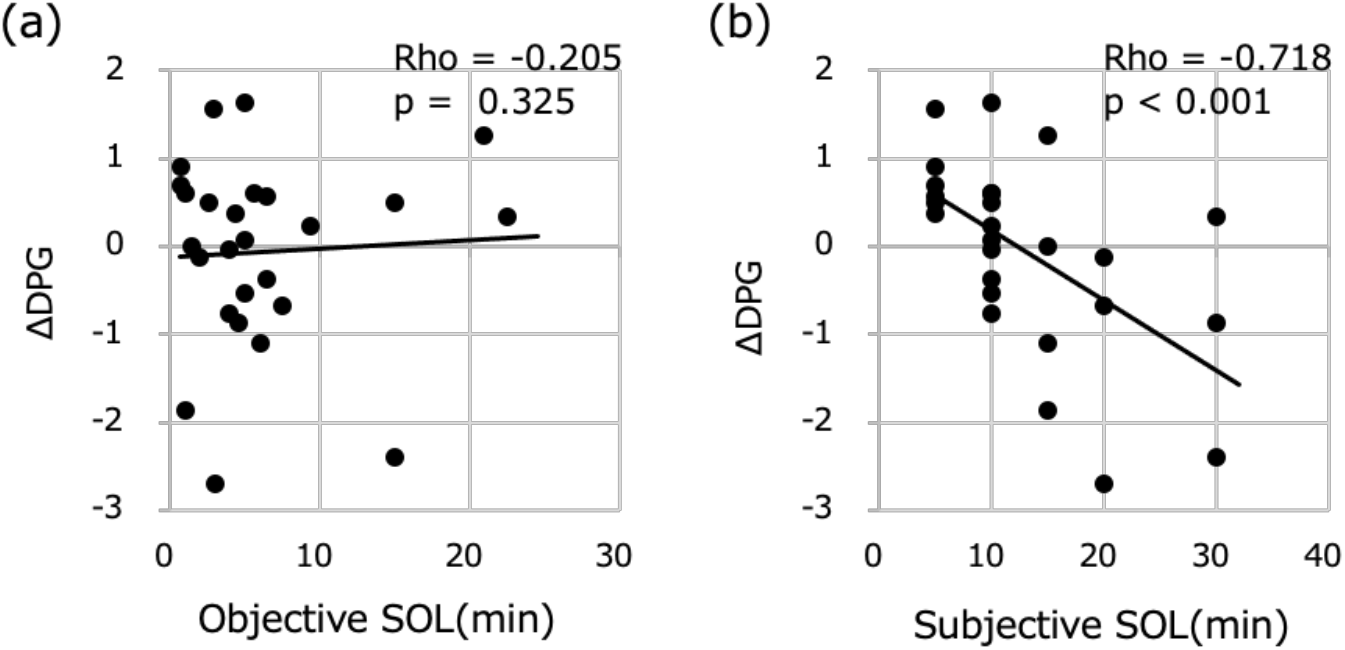
Relationships between mean of DPG during the 15 minutes before the lights-off minus the mean of DPG during the 15 minutes after the lights-off (ΔDPG) and (a) objective SOL, (b) subjective SOL.

### Relationship between subjective SOL and other subjective parameters

Subjective SOL showed significant negative correlations with the subjective sleep depth (Rho = −0.429, p = 0.029), restorative sleepiness (Rho = −0.390, p = 0.045), and total sleep time (Rho = −0.435, p = 0.030), and significant positive correlations with subjective total awaking time (Rho = 0.415, p = 0.039).

### Factors that most influence subjective SOL

Stepwise regression analysis was performed with subjective SOL as the dependent variable and Stage N2 duration, ΔDPG, SWA during sleep in the 15 min after lights-off, subjective sleep depth, and subjective total sleep time, which showed a significant correlation, as independent variables to identify factors that would explain the length of subjective SOL. ΔDPG (p > 0.001, β = 0.574) and stage N2 duration (p = 0.035, β = −0.340) were associated with subjective SOL (0.709, r^2^ = 0.503) (Figure 6). Therefore, ΔDPG was the strongest predictive factor in explaining the length of subjective SOL.

**Fig 6.**
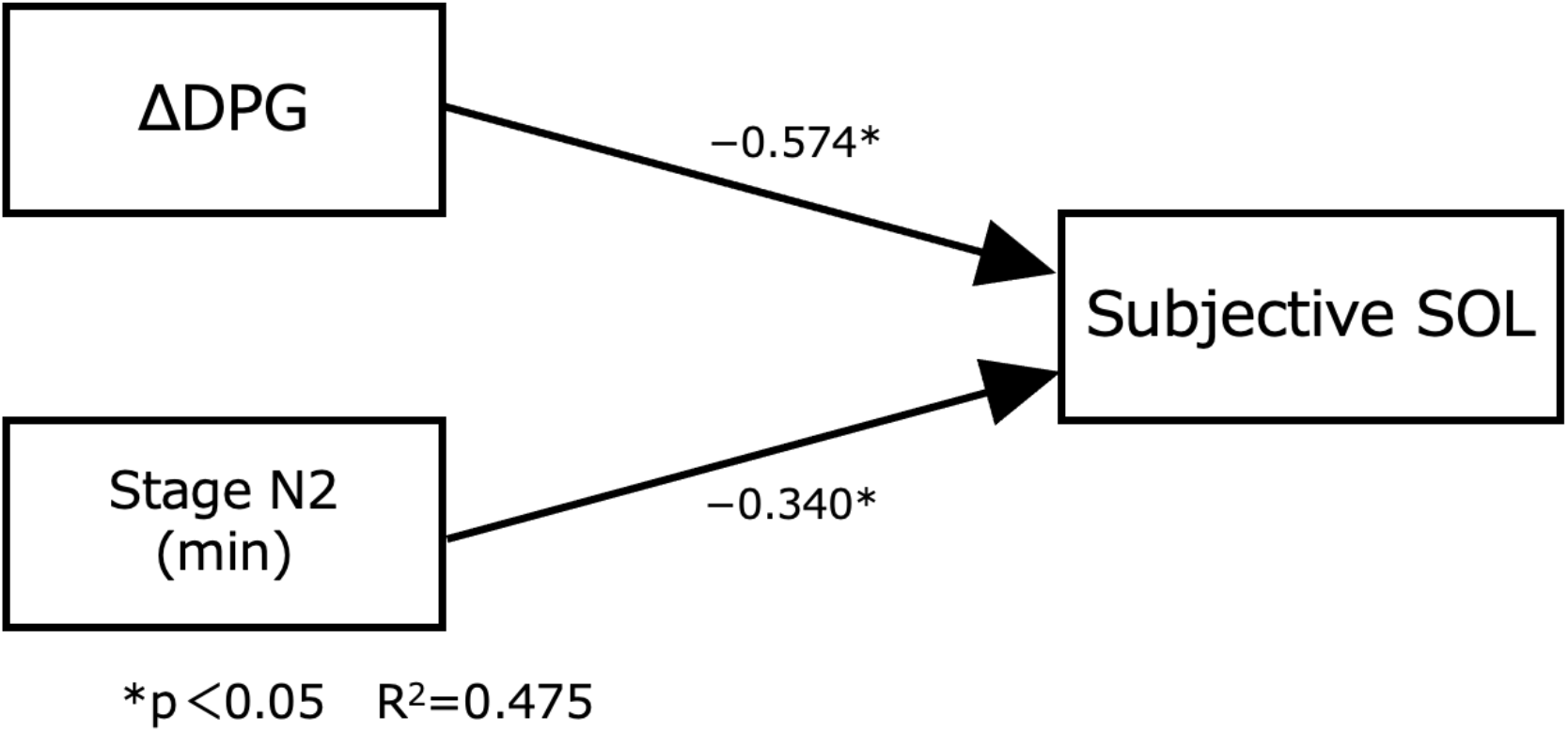
ΔDPG was the most influential factor on subjective SOL, followed by Stage N2 time.

## DISCUSSION

To the best of our knowledge, this is the first study to examine the relationship between the characteristics of subjective SOL, sleep structure, body temperature (skin temperature and heat loss) and subjective evaluation during sleep onset in healthy young adults from the perspective of TEA. The results showed that subjective SOL was negatively correlated with stage N2 duration, SWA, slow wave components immediately after falling asleep, and ΔDPG. Regarding the subjective evaluation of sleep, the results showed significant negative correlations with the subjective ratings of depth of sleep, restorative sleepiness and total sleep time, and significant positive correlations with subjective total awaking time. Furthermore, a stepwise multiple regression analysis revealed that the strongest predictive factor for subjective SOL was ΔDPG, followed by stage N2 duration.

### Subjective and objective SOL

In the present study, most participants evaluated their SOL as being longer than the actual SOL. A study of normal elderly participants in their 60s reported that the latency of falling asleep was subjectively overestimated (subjective SOL of 35.0 min vs. objective SOL of 19.0 min) by approximately 25 min [12]. In addition, several previous studies reported that patients with sleep state misperception significantly underestimated their sleep duration and overestimated SOL [10–12]. Herman et al. investigated people with insomnia aged 50–75 years and reported a discrepancy in the evaluation of SOL of approximately 50 min [12], suggesting a marked difference compared with the healthy participants in the present study. Objective SOL during the daytime for healthy adults is typically approximately 10 min [34], and objective SOL of < 8 min is considered to be diagnostic of excessive sleepiness [35]. In the present study mean SOL of the participants was slightly less than 8 min. This may not be because of excessive sleepiness, but chronic sleep deprivation caused by schoolwork and other factors.

### Subjective SOL and sleep structure

Subjective SOL showed a significant negative correlation with stage N2 duration. This result suggests that the feeling that one has fallen asleep occurs in stage N2, which is a more stable sleep stage. Furthermore, subjective SOL showed a significant negative correlation with slow wave components such as SWA and δ power during the 15 min after lights-off. Previous studies reported that the longer the amounts of SWS that was robust in the first half of the nocturnal sleep period, the longer the participants’ estimates of the subjective sleep duration [1,2]. The present study focused on subjective SOL, and found that the larger the amounts of slow wave components in the sleep onset period after lights-off, the shorter the subjective SOL (i.e., participants fell asleep earlier). The findings of these previous studies and the current results suggest that the slow wave component during the sleep onset period may affect not only subjective sleep duration but also subjective SOL. Despite the possibility that the slow wave component may affect subjective SOL, it was associated with stage N2 duration, not stage N3 duration.

This phenomenon might be caused by the following three factors: 1) K complex is similar to biphasic slow waves with a duration of 0.5 s or longer, 2) delta sleep of less than 6 s in an epoch (30 s) is considered to be stage N2 [32], and 3) the amount of stage N3 was relatively small and varied because participants slept during the daytime period. These factors may have led to the significant negative correlation between subjective SOL and SWA, and the delta sleep component in the present study.

### Subjective SOL and heat loss

The degree of heat loss indicated by DPG is closely related to subjective SOL, and its relationship with objective SOL has been previously reported [22,23]. In accord with this relationship, high DPG (heat loss) before the onset of sleep is caused by an increase in skin temperature and a decrease in CBT, and shortens sleep latency. In the current study, we focused on subjective SOL not objective SOL and examined the relationship with heat dissipation. The results revealed that subjective SOL exhibited a significant negative correlation with ΔDPG that indicated the degree of reduction of heat loss. Therefore, the present results suggest that accelerated heat dissipation (i.e., heat loss) from before lights-off compared with during sleep may enhance the feeling of having fallen asleep earlier.

Kuriyama et al. reported that TEA exhibits diurnal variation and is strongly influenced by changes in body temperature [36,37]. Their study focused on TEA while awake, and they did not examine TEA during sleep. Our previous study also revealed a significant correlation between subjective sleep time and body temperature rhythm [38]. Therefore, subjective SOL may be related to thermoregulatory functions such as heat dissipation and body temperature rhythms. Further studies will be needed to elucidate the detailed mechanisms underlying this phenomenon. In addition, further understanding of the relationship between subjective SOL and heat dissipation in older people, whose thermoregulation functioning is often impaired, would be helpful, as the current study was conducted on healthy young adults.

The current study involved several limitations that should be considered. First, we examined the association between subjective SOL and sleep parameters during daytime sleep, and not during nighttime sleep. Because the phases of sleep structure and body temperature rhythm differ between daytime and nighttime sleep, it is necessary to also study nighttime sleep. Furthermore, PSG measurement was terminated at 60 min after lights-off, suggesting that the difference in sleep stage just before lights-on may affect the subjective evaluation (sleep inertia) after lights-on.

## CONCLUSIONS

The degree of heat dissipation before falling asleep contributed most strongly to the sensation of falling asleep in healthy young adults, and stage N2 duration and slow wave components immediately after falling asleep may be related to the sensation of falling asleep. In addition, when participants experienced a short subjective SOL, they felt that they have had a deeper and longer sleep, were less aware of their total waking time, and were more likely to feel restorative sleepiness. In future, it will be necessary to examine subjective SOL and related factors during nocturnal sleep in patients with insomnia, as well as the effect of heat dissipation on subjective SOL by thermal stimulation, such as exercise, which is believed to promote sleep onset. Examining these issues will be helpful for elucidating the physiological mechanisms of insomnia and its treatment.

## ACKNOWLEDGMENTS

This study was supported by a Grant-in-Aid for Scientific Research from the Japan Society for the Promotion of Science (15K18980). We thank Associate Professor Kozue Kishii of the Graduate School of Saitama Prefectural University for reviewing the paper from various perspectives and providing valuable guidance. We thank Benjamin Knight, MSc., from Edanz (https://jp.edanz.com/ac) for editing a draft of this manuscript.

## AUTHOR CONTRIBUTIONS

R.I., A.K., K.S., M.F., M.H and S.A.-O. conceived and designed research; R.I., A.K., K.S., M.F., M.H and S.A.-O. performed experiments; R.I., A.K., K.S., M.F., M.H and S.A.-O. analyzed data; R.I., M.F. and S.A.-O. interpreted results of experiments; R.I. prepared figures; R.I. drafted manuscript; R.I. and S.A.-O. edited and revised manuscript; R.I., A.K., K.S., M.F., M.H and S.A.-O. approved final version of manuscript.

